# High-throughput, unbiased single-molecule displacement mapping with deep learning reveals spatiotemporal heterogeneities in intracellular diffusivity

**DOI:** 10.1101/2025.02.15.638418

**Authors:** Jiankai Xia, Yi He, Zhipeng Zhang, Jinhong Yan, Ruirong Wang, Mingxuan Tang, Jingye Chen, Jun Fan, Kun Chen

**Affiliations:** School of Optoelectronic Science and Engineering, University of Electronic Science and Technology of China, Chengdu 610054, China; Institute of Fundamental and Frontier Sciences, University of Electronic Science and Technology of China, Chengdu 610054, China; Yangtze Delta Region Institute (Huzhou), University of Electronic Science and Technology of China, Huzhou 313000, China

## Abstract

Single-molecule displacement/diffusivity mapping (SM*d*M) has revolutionized the study of molecular motion in live cells by providing high spatial resolution insights. However, its utility is restricted by measurement biases and low throughput, making it challenging to capture temporal dynamics of intracellular diffusivity. Here we report a high-throughput single-molecule diffusivity microscopy approach (Hi-SM*d*M) that rapidly and unbiasedly maps nanoscale heterogeneities in molecular motion inside mammalian cells, enabling time-resolved imaging of local diffusivity dynamics. Hi-SM*d*M employs a self-supervised deep-learning denoising framework with a general noise model to effectively restore the scarce signals of fast-moving single molecules, while eliminating artifact motions without relying on spatial redundancies or temporal correlations. It then provides unbiased estimations of nanoscale diffusion coefficient and improve both throughput and temporal resolution by up to an order of magnitude. We demonstrate the versatility of Hi-SM*d*M through time-resolved mapping of the spatially heterogeneous diffusivity of free proteins in the cytoplasm. Hi-SM*d*M also unveils the temporal dynamics of intracellular diffusion under hypotonic conditions in live cells. Additionally, Hi-SM*d*M tracks the spatiotemporal dynamics of intraorganellar diffusivity during rapid rearrangements of the ER network and functional activities of mitochondria. Overall, Hi-SM*d*M provides exceptional opportunities for high-throughput and unbiased single-molecule diffusivity mapping with excellent spatiotemporal resolution.

## Introduction

Molecular diffusion in live cells underlies essential biological processes, with the diffusion coefficients *D* of proteins providing valuable insights into cellular crowding, binding interactions, confinement, and microenvironmental effects^1–4^. Such measurements of *D* inform the organization of subcellular structures and the cellular responses to stressors, like osmotic shifts or energy depletion^5, 6^. However, characterizing molecular diffusion in live cells remains challenging due to the spatial and temporal limitations of optical imaging and analysis methods, particularly for freely diffusing, unbound molecules. Conventional fluorescence methods, such as fluorescence recovery after photobleaching (FRAP)^7, 8^ and fluorescence correlation spectroscopy (FCS)^9–11^, estimate *D* from ensembles of molecules but offer limited capabilities for spatial mapping. Single-molecule tracking (SMT) excels in following long trajectories of membrane-bound, slow-diffusing molecules, especially within small bacterial volumes^12–14^. Yet, applying SMT to freely diffusing molecules in eukaryotic cells proves difficult, as these molecules rapidly diffuse out of the focal plane, preventing tracking over many consecutive frames.

To address this, we recently developed single-molecule displacement/diffusivity mapping, named SM*d*M, echoing the concept of single-molecule localization microscopy (SMLM)^15^, which detects the transient displacement of single molecules across tandem camera frames under synchronized stroboscopic illumination^16^. By requiring only two localizations within a short time window, SM*d*M effectively captures the motion of fast, unbound molecules without needing long trajectories. By locally accumulating mass of single-molecule displacements over frames^17^, SM*d*M generates super-resolution maps of diffusion coefficients, revealing nanoscale heterogeneities in diffusivity for both free proteins and small solutes in live mammalian cells^18–20^.

While this location-oriented strategy is naturally powerful for spatial mapping, SM*d*M struggles to capture temporal dynamics in molecular diffusivity^16^. SM*d*M adapts very short excitation pulses (∼0.5 ms) to minimize motion blur from freely-diffusing molecules. However, this stroboscopic illumination approach inevitably reduces the brightness and signal-to-noise ratio (SNR) of fluorescent molecules, thereby limiting the number of detectable molecules and lowering the overall throughput. Furthermore, SM*d*M requires each molecule to be captured in two consecutive frames and correctly paired, yet nearly 50% of detected molecules are mis-paired or unpaired, further reducing effective throughput.Constructing a meaningful super-resolution diffusivity *D* image of cytoplasm demands ∼150,000-200,000 frames (∼22-30 minutes at 110 fps), making real-time intracellular diffusivity monitoring impractical^16, 17^. This prolonged acquisition time also introduces substantial phototoxicity, further limiting live-cell imaging. Additionally, SM*d*M exhibits inherent detection bias: brighter, slower-moving molecules are more easily detected, while dimmer, faster-moving molecules disperse widely and often remain undetected. This exclusion skews results toward the slow-diffusion regime.

In this work, we leverage advancements in deep learning to overcome these limitations by implementing a two-stage denoising framework tailored for freely diffusing single-molecule data. This optimized framework enables accurate recovery of valid molecules from noisy single-molecule images without relying on spatial or temporal redundancies or long-range correlations, thereby minimizing potential distortions in molecular diffusivity. We thus can quantify all effective moving-molecules without dependence on their brightness, thereby enabling high-throughput, unbiased single-molecule displacement/diffusivity mapping (Hi-SM*d*M) at the super-resolution level. Hi-SM*d*M unveils rich temporal dynamics of intracellular diffusion in live cells with up to 10-fold improved temporal resolution, along with spatial heterogeneity at the nanoscale.

## Results

### SM*d*M inherently underestimate diffusion coefficients *D* of fast-moving molecules

For SM*d*M of free proteins (mEos3.2) in the cytoplasm of COS-7 cells, we employ paired, stroboscopic excitation pulses across tandem camera frames with 500 μs duration separated by 1 ms to minimize motion blur, enabling the precise localization and tracking of diffusing molecules and yielding their nanoscale displacements (*d*) (Fig. 1a-b). By spatially binning the displacements accumulated over ∼7.5×10⁴ tandem-excitation cycles onto 100 nm grids and fitting them to a random-walk model using maximum likelihood estimation (MLE) (Methods), we achieved super-resolution mapping and quantification of *D* (Fig. 1c). However, our findings reveal an inherent measurement bias SM*d*M. Diffusion coefficient estimations are affected by the brightness of the molecules, as slower molecules appear brighter, while faster ones are dimmer and harder to detect (Fig. 1b). This bias likely underestimates diffusivity in *D* mapping (Fig. 1c), as fast-moving molecules are excluded from displacement calculations. To further examine this effect, we analyzed displacement and brightness across 150,000 frames, revealing an inverse correlation (Fig. 1d). Varying localization thresholds—while maintaining molecule count and reliable fits (Methods)—showed that higher thresholds reduce global diffusivity, while lower ones increase it (Fig. 1e, f; Supplementary Fig. S1). Simulations confirmed this brightness-dependent bias (Supplementary Fig. S2). Reducing this bias requires detecting all molecules regardless of brightness, but practical challenges remain. SM*d*M’s short stroboscopic pulses lower SNR, making fast-moving molecules register near background noise. Lowering thresholds captures some dim, fast molecules but also increases noise, degrading *D* fitting accuracy (Fig. 1g, h).

**Figure 1.**
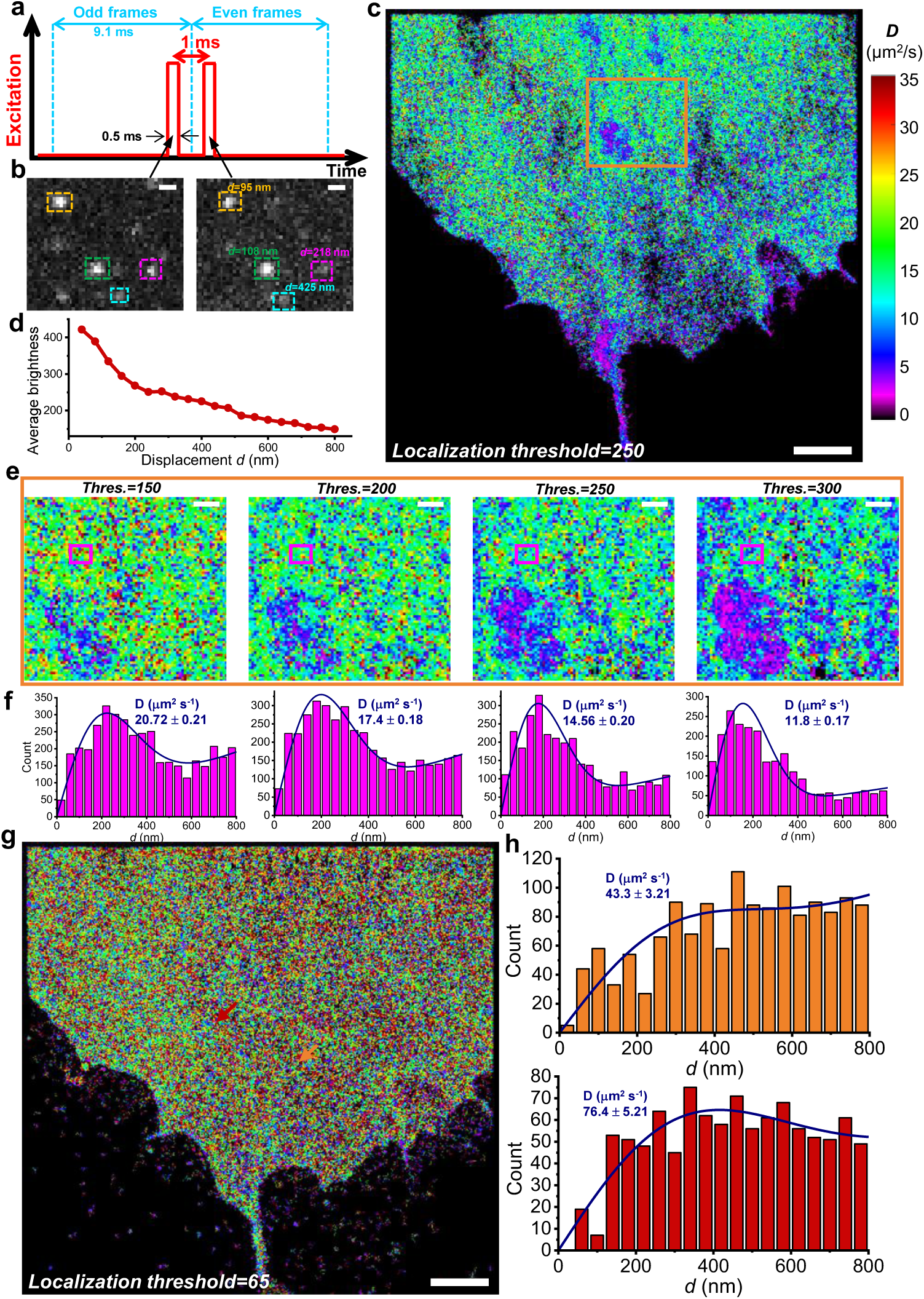
The inherent measurement bias of SM*d*M to underestimate the diffusivity of fast-moving molecules. **a.** Schematics: tandem excitation pulses of *τ* = 500 μs duration are applied across paired camera frames at a Δ*t* = 1 ms center-to-center separation, and this scheme is repeated ∼7.5×10^4^ times to enable local statistics. **b.** Example single-molecule images captured in a tandem frame pair for mEos3.2 diffusing in the cytoplasm of a live COS-7 cell, with matched pairs marked by same-colored boxes, showing varying displacements. Dimmer molecules correspond to larger displacements. **c.** SM*d*M-reconstructed super-resolution *D* map (bin size: 100×100 nm^2^) in COS-7 cytoplasm, accumulated 150,000 frames, with a threshold of 250 for single-molecule localization. **d.** Inverse relationship between the average brightness and displacements of all single molecules across 150,000 frames. **e.** Reconstructed SM*d*M *D* maps for a subregion (orange box in c) under different single-molecule localization thresholds, showing reduced global *D* values with higher thresholds. **f.** Distribution of the SM*d*M-measured 1-ms single-molecule displacement *d* for boxed regions in e (magenta). Blue curves: Fits to the SM*d*M diffusion model, with resultant *D* values marked in each plot. **g.** The reconstructed SM*d*M *D* map for the same cell in c at a significantly low localization threshold shows higher diffusion coefficients but increased noise. **h.** Representative displacement distributions for two 300×300-nm^2^ areas in g (red and orange arrows) highlight poor fits due to excessive noise. Scale bars: 1 μm (b, e); 5 μm (c, g).

### Construction of deep-learning denoising framework for fast-diffusing molecule Imaging

Accurate restoration of fast-diffusing single molecules, regardless of noise, is essential for achieving high-throughput, unbiased single-molecule diffusivity mapping. The rapid evolution of deep learning has notably impacted biological fluorescence imaging^21^, especially in denoising applications for photon-limited conditions, such as single-molecule microscopy^22–25^. In practice, self-supervised deep-learning denoising has increasingly supplanted supervised methods due to the limitations of experimental imaging conditions^26, 27^. Currently, most self-supervised approaches in fluorescence imaging rely on Noise2Noise-like frameworks, utilizing either spatial redundancy from neighboring pixels or temporal redundancy from slowly moving cellular structures^28–30^. However, single-molecule imaging lacks temporal redundancy due to the random blinking of individual molecules. To address this, the latest SRDTrans algorithm enhances denoising by leveraging spatial redundancy and long-range temporal correlations, optimized for SMLM of fixed cells^31^. This approach has achieved state-of-the-art performance in denoising single-molecule images in super-resolution microscopy. However, for freely diffusing single molecules, both random blinking and movement occur on short timescales. While temporal correlation aids in restoring static single-molecule blinking, it cannot accurately estimate spatial displacement due to molecular motion. Instead, it naturally interprets moving molecules as stationary, underestimating metrics such as displacement and diffusivity.

To overcome this limitation, we present a denoising framework for SM*d*M (Hi-SM*d*M) that leverages an alternative approach: denoising through noise modeling^32^. While noise modeling is a common technique in computer vision^33–37^, it is typically applied in a supervised context. In contrast, fluorescence images captured directly by scientific cameras—without in-camera signal processing pipeline^38^—retain two valuable properties: strong spatial correlation between neighboring pixels and independent noise components. These features enable the use of self-supervised noise modeling for denoising, which is particularly suited for single-molecule imaging. Our network, outlined in Fig. 2a, begins with the synthesis of a clean single-molecule image. A corresponding low-SNR image is then generated using noise modeling, and this paired data is used to train a denoising network^39^ (Fig. 2b). Once trained, the network can be applied to denoise real noisy images. The key noise modeling process operates on three levels: global noise level (𝐺), local noise map (𝑀), and pixel-wise noise value (𝑉). By applying adjacent random masking, we estimate approximate clean signal values from the center pixel and its neighboring pixels and calculate 𝐺, 𝑀, and 𝑉 from them for corresponding noise (methods detailed in Supplementary Note 1). To ensure noise independence, three separate, non-overlapping random samplings are taken from the neighborhood when computing 𝐺, 𝑀, and 𝑉. The nearly clean signal mask is obtained by inverting these random masks within the neighborhood. At the network level, we designed three distinct modules: the Global Noise Level Predict Module to predict *Ĝ*; the Local Noise Map Predict Module to incorporate *Ĝ* and predict *M̂*; and the sampling module to generate *V̂* based on *M̂*. Loss functions across 𝐺, 𝑀, and 𝑉 are used to iteratively update the noise modeling network (Fig. 2c, Supplementary Note 1).

**Figure 2.**
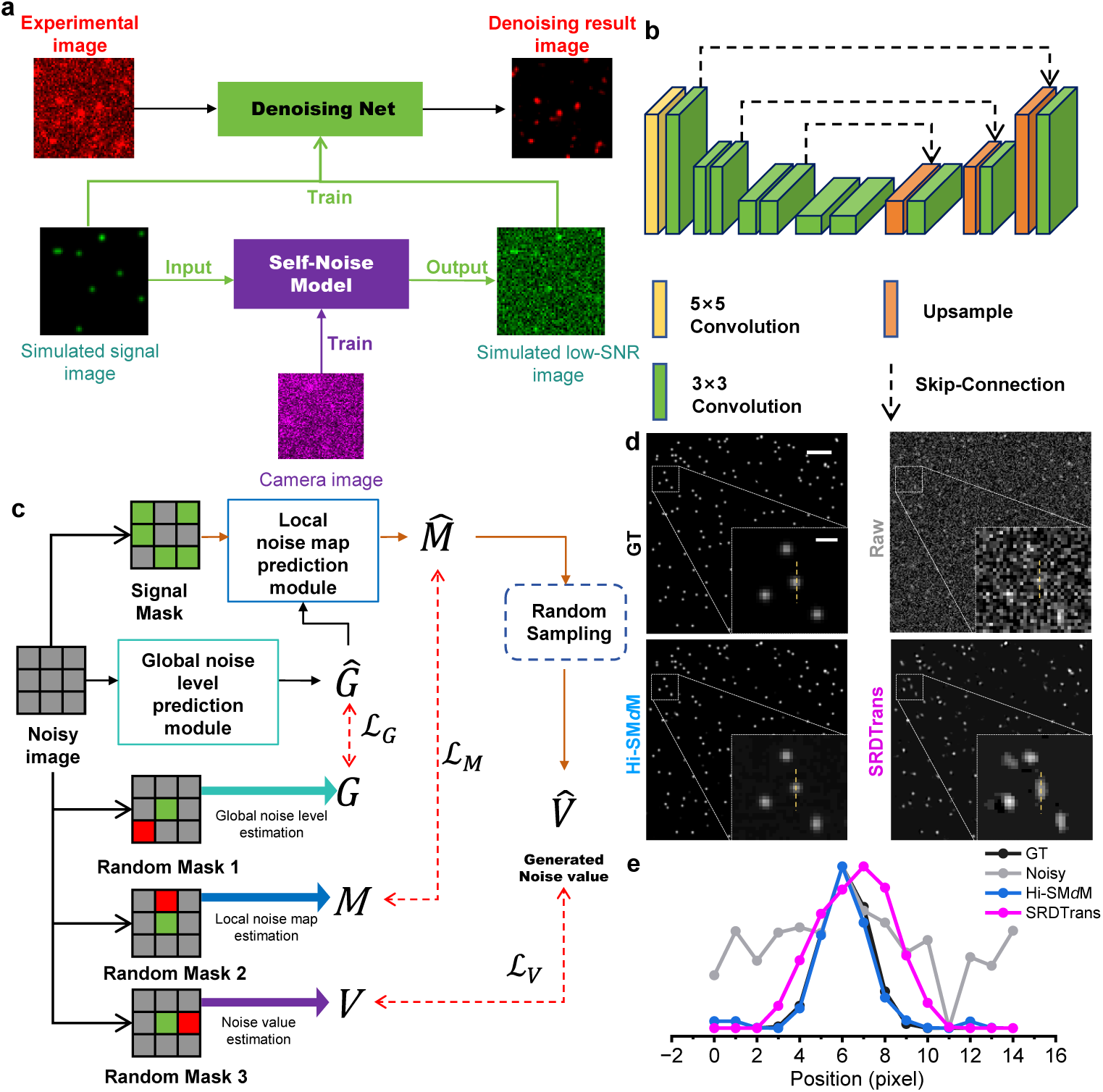
Principle of Hi-SM*d*M framework and denoising performance evaluation. **a**. Blind denoising scheme. A self-supervised noise model is trained using images captured by a scientific camera. Clear single-molecule images are simulated using a point spread function combined with 2D Brownian random walks, with intensity levels matching experimental data. The model generates paired low-SNR images corresponding to the simulated signal images, which are used to train a denoising network. The trained network is then applied to experimental data, producing denoised images. **b**. Denoising network architecture. The network utilizes a U-Net structure with convolutional layers and skip connections. Low-SNR images serve as inputs, clear images as targets, and the network is optimized with a mean squared error (MSE) loss function. **c**. Structure of self-supervised noise modeling. The noise model includes three modules: global noise level prediction, local noise map prediction, and noise sampling. Noise is approximated using a random mask and modeled at global (*G*), local (*M)*, and pixel (*V*) levels. The loss function is a weighted sum of three individual losses (ℒ_𝐺_, ℒ_𝑀_, ℒ_𝑉_) corresponding to these levels. For details, see Supplementary Note 1. **d**. Performance comparison of Hi-SM*d*M and SRDTrans for single-molecule image denoising. Both methods blindly denoise simulated single-molecule diffusion data, demonstrating the superior performance of Hi-SM*d*M. **e**. Intensity profiles along the yellow dashed lines in d, highlighting the effectiveness of Hi-SM*d*M in preserving signal integrity. Scale bar: 5 μm and 1 μm for magnified view (d) .

To demonstrate the effectiveness of our denoising network (Hi-SM*d*M), we generated simulated diffusing single-molecule data by randomly selecting the starting positions for molecules with a typical cytoplasmic diffusion coefficient of 20 μm^2^/s at every micro-step. Molecules were programmed to disappear after diffusing for a random duration, simulating either movement out of the field of view or photobleaching, as seen in real experiments. The simulations were conducted under typical experimental conditions, with a pixel size of 160 nm and an exposure time of 0.5 ms per pulse, with the two tandem pulses separated by 1 ms. To generate noisy data, a high level of mixed Poisson–Gaussian noise was applied to the noise-free single-molecule images, and the resulting noisy images were used as input for the denoising schemes (Fig. 2d). In the raw images, few molecules were correctly detected. Although both Hi-SM*d*M and SRDTrans can generate denoised image of diffusing molecules with substantially improved SNR, quantitative comparisons of the visualized images (Fig. 2d and Supplementary Video 1) and the extracted intensity profiles (Fig. 2e) show that Hi-SM*d*M aligns closely with the ground truth (GT). In contrast, SRDTrans produces elongated and blurred features in single-molecule images due to its reliance on long-range dependencies across multiple frames—an issue that does not impact fixed molecules in conventional SMLM.

### Performance characterization of Hi-SM*d*M for spatial diffusivity mapping with simulated data

We next examined how denoising affects the extracted step displacements (*d*) of diffusing molecules between tandem frames. By merging odd and even frames, each molecule pair became clearly visible, allowing direct assessment of step displacements (Fig. 3a). In raw, noisy images, identifying molecule pairs was challenging, while Hi-SM*d*M clearly preserved molecule pairs consistent with the GT. In contrast, SRDTrans introduced artificial proximity between paired molecules, leading to increased spatial overlap (highlighted in yellow) and displacement shortening (see zoom-in regions and intensity profiles in Fig. 3a). This artifact is quantified in Fig. 3b, where SRDTrans’s elongated single-molecule features result in reduced step displacements *d*, while Hi-SM*d*M accurately preserves single-molecule diffusivity. The robust denoising provided by Hi-SM*d*M significantly enhances detection, recovering nearly all valid molecules and increasing localization throughput by 4-fold and molecule pairing by 8-fold compared to raw data (Fig. 3c).

**Figure 3.**
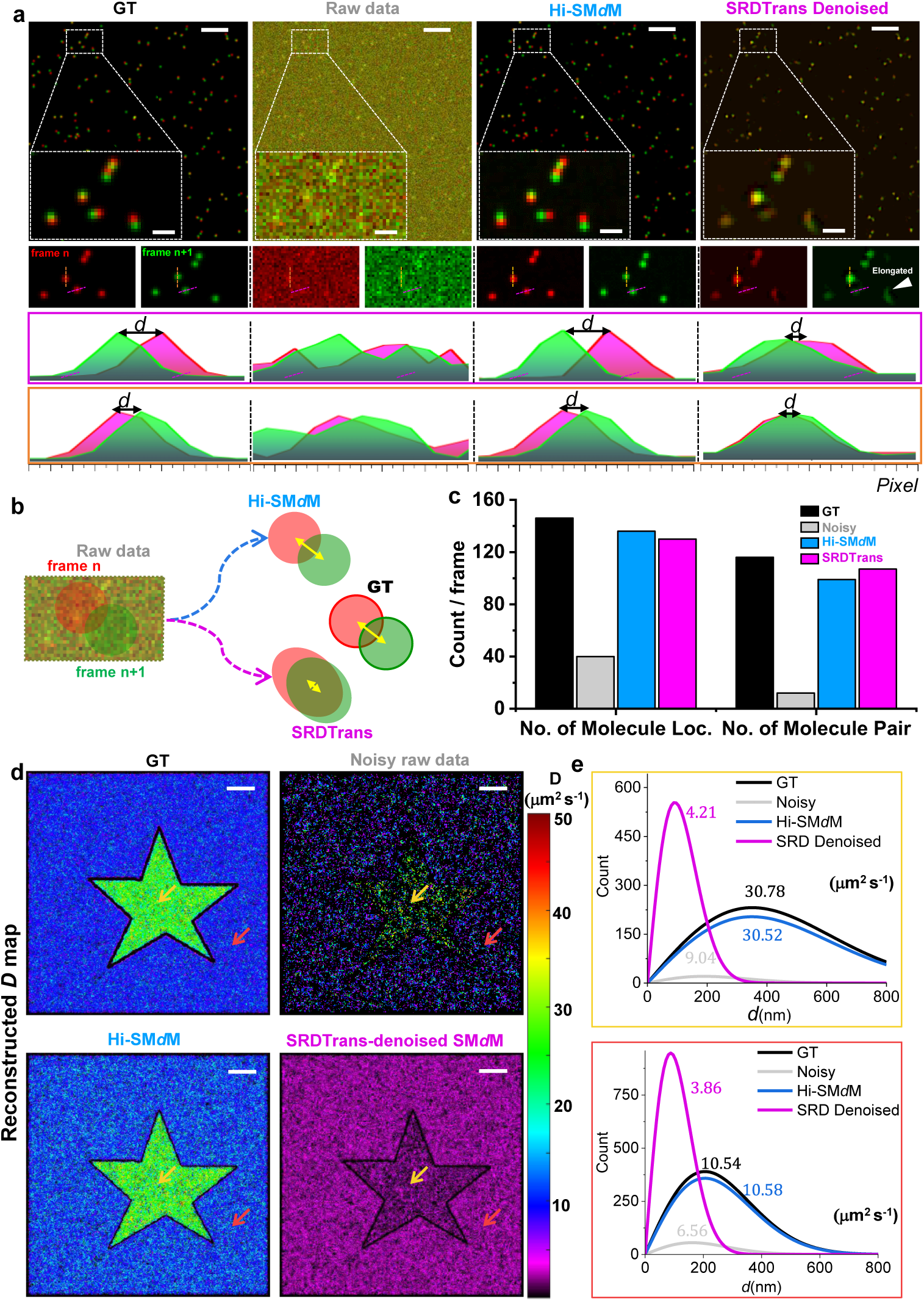
Performance assessment of diffusivity mapping using Hi-SM*d*M on simulated single-molecule diffusion data. **a.** Visualization of tandem frames for the ground truth (GT), raw noisy images, Hi-SM*d*M, and SRDTrans-denoised images. Zoom-in boxes highlight paired diffusing molecules. Bottom: Extracted intensity profiles for two molecule pairs (magenta and orange dashed lines). Hi-SM*d*M effectively preserves molecule pair identity, consistent with the GT. In contrast, SRDTrans results in increased spatial overlap and shortened displacements. **b.** Schematic illustration of the effects of Hi-SM*d*M and SRDTrans denoising on single-molecule displacement analysis. **c.** Comparison of single-molecule localizations and matched molecule pairs per frame for Hi-SM*d*M and SRDTrans, compared to Ground Truth (GT) and raw noisy images. Both methods recover nearly all valid molecules, increasing localization throughput by 4-fold and molecule pairing by 8-fold compared to raw data. **d.** Reconstructed diffusivity (D) maps using GT, raw noisy images, Hi-SM*d*M-denoised, and SRDTrans-denoised images. Only Hi-SM*d*M reproduces the diffusivity distribution consistent with the GT. **e.** Representative displacement distributions for two 400×400-nm^2^ areas, one inside (top: yellow arrow in d) and one outside (bottom: red arrow in d) the star-shaped region, accumulated from GT, raw noisy images, Hi-SM*d*M-denoised, and SRDTrans-denoised images. Scale bars:5 μm and 1 μm for magnified view (a); 5 μm(d).

To further examine the performances of Hi-SM*d*M for diffusivity mapping, we simulated spatial patterns with distinct diffusion coefficients, such as star-shaped regions with a high *D*-value of 30 μm^2^/s at the center and a surrounding region with a low *D*-value of 10 μm^2^/s. The simulated diffusing molecules over 10,000 frames were localized and spatially binned into a 120 nm × 120 nm grid. By fitting accumulated displacements *d* in each bin through MLE we derived nanoscale diffusion coefficients *D*, allowing us to render a super-resolution map of local diffusivity across the entire field of view (Fig. 3d). We thus showed that Hi-SM*d*M correctly mapped out spatial differences in diffusivity by detecting almost all molecules and preserving their true displacements. In contrast, raw, noisy data lacked sufficient valid molecule localizations, introducing numerous artifacts that obscured meaningful structural information in the *D* map. Although SRDTrans detected a comparable number of molecules, it significantly underestimated the diffusion coefficient *D* across the map, consistent with artifacts that reduce observed step displacements. Quantitative results from two spots in the high- and low-*D* regions (Fig. 3e), for bins 360 × 360 nm^2^ in size, further highlight the high accuracy of Hi-SM*d*M in diffusivity mapping. Together, these simulations demonstrate Hi-SM*d*M’s robust, high-throughput, and unbiased performance for diffusivity mapping.

### Hi-SM*d*M unveils nanoscale changes in diffusivity as cellular organelles undergo dynamic structural rearrangements over time

To evaluate Hi-SM*d*M under challenging conditions, we imaged free mEos3.2 proteins in the cytoplasm of COS-7 cells during time-lapse live-cell experiments. Although mEos3.2 is a widely used photoswitchable fluorescent protein in SMLM^40^, its limited photon budget complicates SM*d*M, especially in COS-7 cells, which have a more three-dimensional morphology and higher background than PtK2 cells used in our previous experiments^16^. These factors severely reduce SNR and throughput, making SM*d*M measurements particularly demanding. Despite this, Hi-SM*d*M effectively restored single-molecule signals and accurately paired molecules between frames (Supplementary Fig. S3 and Video 2).

Using raw images, SM*d*M attempted to reconstruct super-resolved *D*-maps with a bin size of 100 nm × 100 nm. With 25,000 frames, the reconstructed diffusivity mapping was incomplete, leaving large areas within the COS-7 cell unresolved (Fig. 4a (I) and Supplementary Fig. S4). Even with 100,000 frames, gaps persisted due to insufficient displacement data (white arrowheads in Fig. 4a (II)). A complete *D*-map required 200,000 frames, revealing nanoscale diffusivity heterogeneities (Fig. 4a (III)). While the local single-molecule displacement distributions were well-fitted to our model, SM*d*M revealed *D* of 10–15 μm^2^/s in the high-*D* regions, significantly slower than the cytoplasmic diffusion previously observed in PtK2 cells (Fig. 4b)^16^. In contrast, Hi- SM*d*M using denoised images required only 25,000 frames to reconstruct complete and super-resolved *D* maps (Fig. 4c), achieving an 8-fold throughput increase while preserving nanoscale diffusivity heterogeneities (Fig. 4c and 4d). Additionally, it reported higher diffusion rates (20–25 μm^2^/s, Fig. 4c-e), aligning better with prior findings. These results highlight Hi-SM*d*M’s ability to correct biases inherent to SM*d*M, particularly in detecting fast-moving molecules under low-SNR conditions.

**Figure 4.**
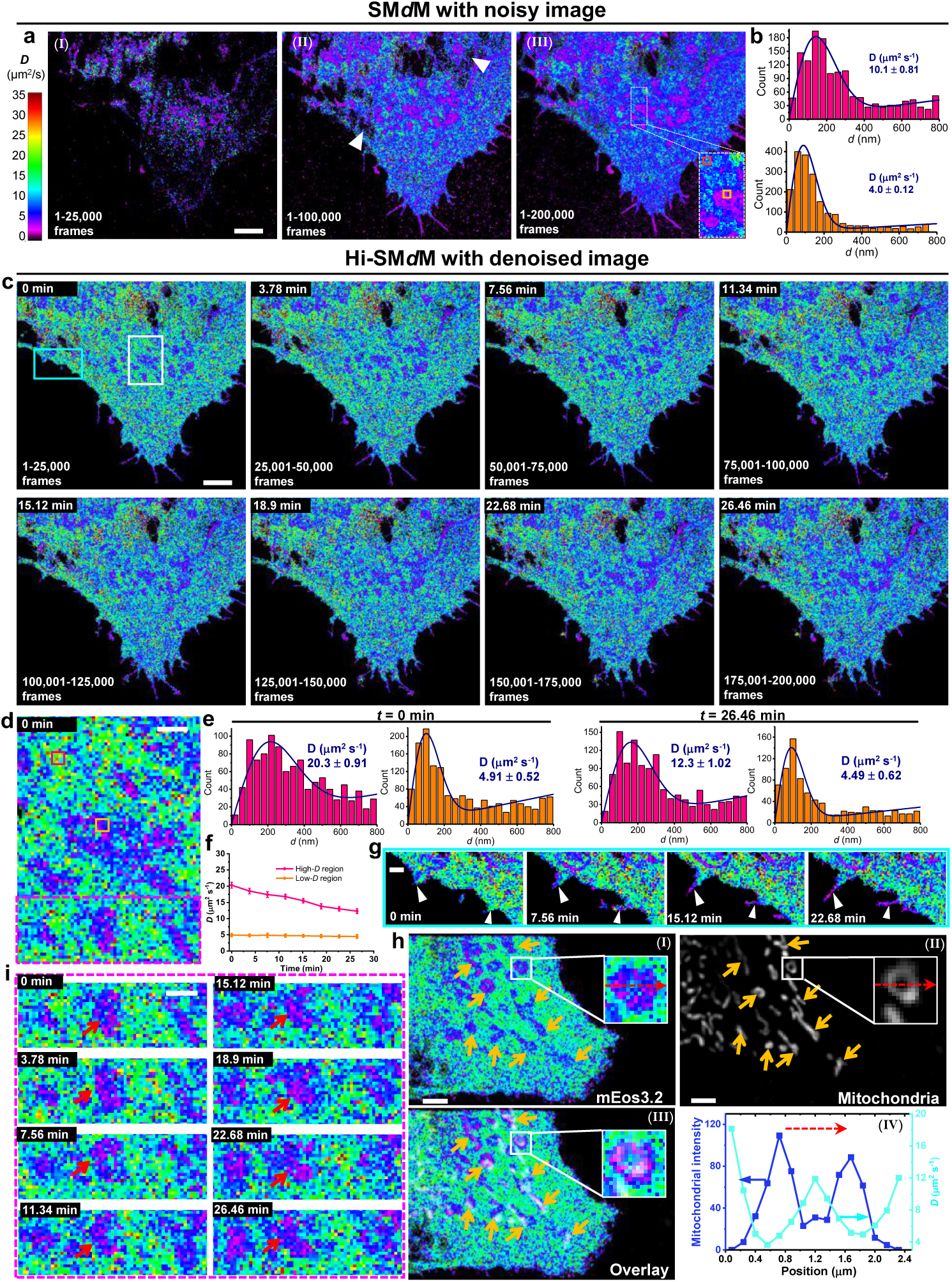
Hi-SM*d*M unveils nanoscale changes over time in cytoplasmic diffusivity of live COS-7 cells. **a.** SM*d*M-reconstructed *D* map (bin size: 100×100 nm^2^) with 25,000 frames (I), 100,000 frames (II), 200,000 frames (III), respectively. **b.** Distribution of single-molecule displacement *d* for boxed regions in a (red and orange). Blue curves: Fits to the diffusion model, with resultant *D* values marked in each plot. **c.** Hi-SM*d*M-reconstructed super-resolution *D* map (bin size: 100×100 nm^2^) for every 25,000 frames, thus enables the time-resolved imaging of diffusivity with a moderate time interval of 3.78 minutes. **d.** Zoomed-in view of a region in (c) (white box), highlighting distinct high- and low-*D* regions in the cytoplasm. **e.** Displacement distributions for two 1 × 1 mm^2^ areas in the high- (red box) and low-*D* regions (orange box) in d at *t* =0 min and *t* =26.46 min. **f.** The variation of *D* with time in the same high- (red box) and low-*D* regions (orange box) as e. **g.** Zoomed-in view of a region in (c) (cyan box) showing the growth of apoptotic cilia over time, with notably low diffusivity in these confined structures. **h.** Another example of Hi-SM*d*M diffusivity map of mEos3.2 in the cytoplasm of a live COS-7 cell (I). Conventional fluorescence image of the same fixed cell using the mitochondria stain MitoTracker Deep Red FM (II). Overlay of I and II (III). Variation in the *D* (cyan line) and the mitochondrial intensity (blue line) along the horizontal direction for a donut-like mitochondrion marked by the red arrows in I and II. **i.** Time-resolved imaging of *D* in the boxed subregion in d (magenta) at 3.78-min intervals, showing the dynamics of mitochondrion-induced low-*D* regions. Scale bars: 5 μm(a, c, h); 1 μm (d, g, i, h).

With its high-throughput and unbiased single-molecule diffusivity mapping capability, Hi-SM*d*M enables, for the first time, investigation of the temporal dynamics of cellular diffusivity at high spatial resolution (Fig. 4c and Supplementary Video 3). Over extended imaging periods, we thus visualized a gradual decrease in cytoplasmic diffusivity (Fig. 4c and another example in Supplementary Fig. S5). Quantitative analysis of high- and low-*D* regions (Fig. 4e and 4f), for bins 1 × 1 μm^2^ in size (red and orange boxes in Fig. 4d), further verified the significant reduction of *D* in the faster-moving regions. This decrease is likely due to cell apoptosis induced by prolonged laser illumination, which increases intracellular viscosity^41^. The unhealthy state of the cells was further evidenced by the growth of apoptotic cilia over time, where diffusivity was notably low in these confined structures (Fig. 4g). Furthermore, Hi-SM*d*M revealed substantial low-*D* regions in the cytoplasm of COS-7 cell (∼5 μm^2^/s, Fig. 4c and 4d) exhibiting distinct characteristics compared to the linear diffusivity patterns in PtK2 cells, which were previously attributed to actin bundles^16^. To investigate the origin of these heterogeneities, we first performed Hi-SM*d*M imaging of mEos3.2 in another live COS-7 cell, followed by concurrent imaging of mitochondria using MitoTracker Deep Red FM in the same cell (Fig. 4h). The low-*D* regions closely corresponded to mitochondrial structures (Fig. 4h and Supplementary Fig. S6). Plotting the measured *D* values across a circular hollow mitochondrion (zoom-in box in Fig. 4h) revealed an inverse relationship with mitochondrial intensity (Fig. 4h (IV)), where mitochondria imped intracellular diffusivity and faster diffusion was observed in the hollow center. This finding highlights the excellent spatial resolution of Hi-SM*d*M. Hi-SM*d*M further refined the full width at half maximum (FWHM) of a tubular mitochondrion from 540 nm (diffraction-limited) to 380 nm (Supplementary Fig. S6), confirming its super-resolution capability. These results underscore the power of Hi-SM*d*M in resolving intracellular diffusion dynamics at sub-diffraction spatial scales.

Hi-SM*d*M *D* maps thus revealed the dynamic structural rearrangements and movements of mitochondria over time (red arrows in Fig. 4i and Supplementary Video 4) by tracking the temporal changes in diffusivity in the slowdown regions that SM*d*M could only resolve as time-averaged results. Unlike in PtK2 cells, COS-7 cells lacked actin-induced slowdown, likely due to their rounded morphology. These findings highlight the critical role of nanoscale structures and organelles in shaping intracellular diffusion.

### Hi-SM*d*M unveils spatiotemporal dynamics of intracellular diffusivity under hypotonic conditions in live cells

Macromolecular crowding plays a crucial role in regulating protein activities and functions within cells, and molecular diffusion offers a direct assessment of this crowding^16, 42^. In this study, we used Hi-SM*d*M to investigate temporal changes in nanoscale crowding in live cells subjected to 50% osmotic pressure. Super-resolution diffusivity maps, reconstructed from denoised images, achieved a temporal resolution of 3.78 minutes, revealing substantial spatiotemporal heterogeneities in macromolecular crowding in live cells.

In untreated COS-7 cells, cytoplasmic diffusivity was largely uniform (Fig. 5) or highlighted heterogeneity in slow-diffusion regions associated with mitochondria (Supplementary Fig. S7). Upon 50% hypotonic treatment, cytoplasmic diffusion initially increased by ∼20% over ∼10 minutes before gradually decreasing (Fig. 5a, 5b and Supplementary Video 4), overshooting to a value ∼16 μm^2^/s lower than the starting *D*, and partially recovering (Fig. 5a, 5b and Supplementary Video 4). These temporal changes indicate osmotic stress-induced variations in cytoplasmic crowding. The findings suggest that hypotonic conditions cause cell swelling, leading to a reduction in macromolecular crowding, followed by recovery through regulatory volume increase mechanisms^43^. This gradual expansion and recovery are consistent with previous results using a FRET-based crowding sensor^44^ (Supplementary Fig. S8). Notably, we observed variability in cellular responses, with diffusion rates in some cells failing to return to baseline levels after the initial increase (Supplementary Fig. S7). A similar behavior was also observed in our cytoplasmic crowding experiments conducted with the FRET sensor with the excitation spectral microscopy^45^ (Supplementary Fig. S8). At the subcellular level, however, diffusion coefficients within mitochondrial regions remained largely unchanged under the same treatment (Supplementary Fig. S7). However, unlike Hi-SM*d*M, bulk-averaged FRET measurements could not resolve nanoscale crowding heterogeneities. Since FRET crowding sensors are excluded from mitochondria, they miss the significantly reduced diffusivity in these compartments. The cellular interior is inherently heterogeneous, consisting of distinct compartments with unique properties. Hi-SM*d*M uniquely integrates super-resolution imaging with single-molecule diffusion analysis, providing a powerful approach for uncovering nanoscale crowding dynamics and their functional implications across distinct cellular structures.

**Figure 5.**
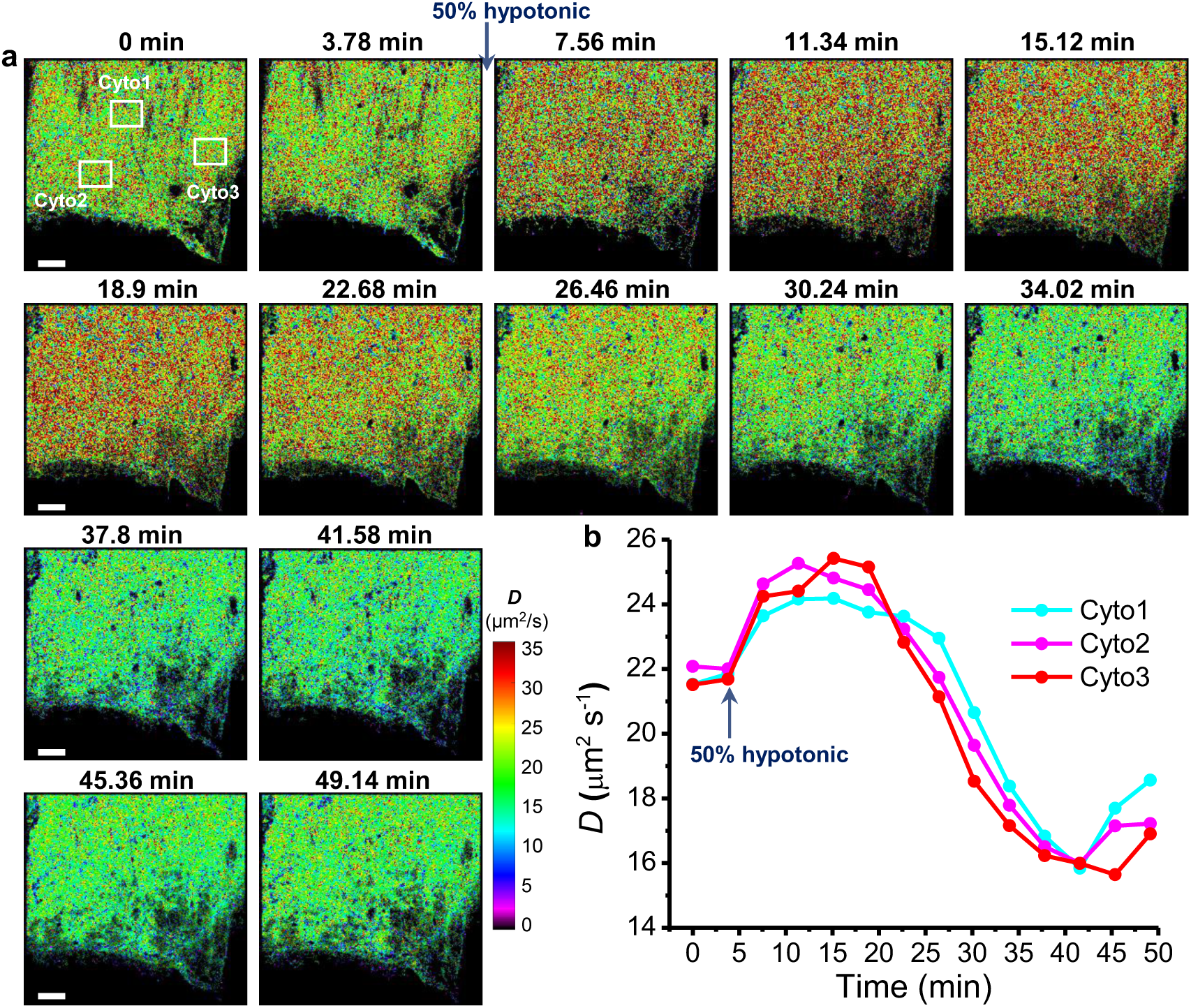
Hi-SM*d*M unveils temporal dynamics of intracellular diffusivity under hypotonic conditions in live cells. **a.** Time-resolved imaging of full *D* maps with Hi-SM*d*M in cytoplasm of a live COS-7 cell at 3.78-min intervals. A 50% hypotonic treatment was applied by adding an equal volume of water to the medium at *t* =7.56 min. Bin size: 160×160 nm^2^. **b.** Temporal variation of *D* in the boxed regions in a, highlighting changes in intracellular diffusivity during hypotonic treatment. Scale bars: 5 μm (a).

### Hi-SM*d*M unveils spatiotemporal dynamics of intraorganellar diffusivity in ER

To further challenge the spatiotemporal resolution of Hi-SM*d*M, we applied it to the highly dynamic endoplasmic reticulum (ER), a continuously rearranging network of tubules and sheets. Previously, resolving diffusion within the ER lumen required ∼100,000 frames for analysis^19^. Using the same photoswitchable fluorescent protein (Dendra2) in the ER lumen, Hi-SM*d*M successfully reconstructed the intricate ER network with an order of magnitude fewer frames (Fig. 6a). By contrast, while SM*d*M could also generate a complete ER diffusivity map after accumulating 100,000 frames, the extended imaging duration (∼15 minutes) caused significant motion blur, obscuring the fine details of the ER tubule network (Fig. 6b). As noted in previous studies, resolving the delicate morphology of ER tubules is challenging with SM*d*M due to its limited spatial and temporal resolution. Additionally, SM*d*M revealed a lower average diffusion rate (7.5 μm^2^/s in the red box, Fig. 6b) compared to Hi-SM*d*M (10.7 μm^2^/s in the white box, Fig. 6a), addressing the bias of SM*d*M against faster-moving molecules. Moreover, SM*d*M failed to produce a high-resolution diffusivity map with only 10,000 frames (Fig. 6c).

**Figure 6.**
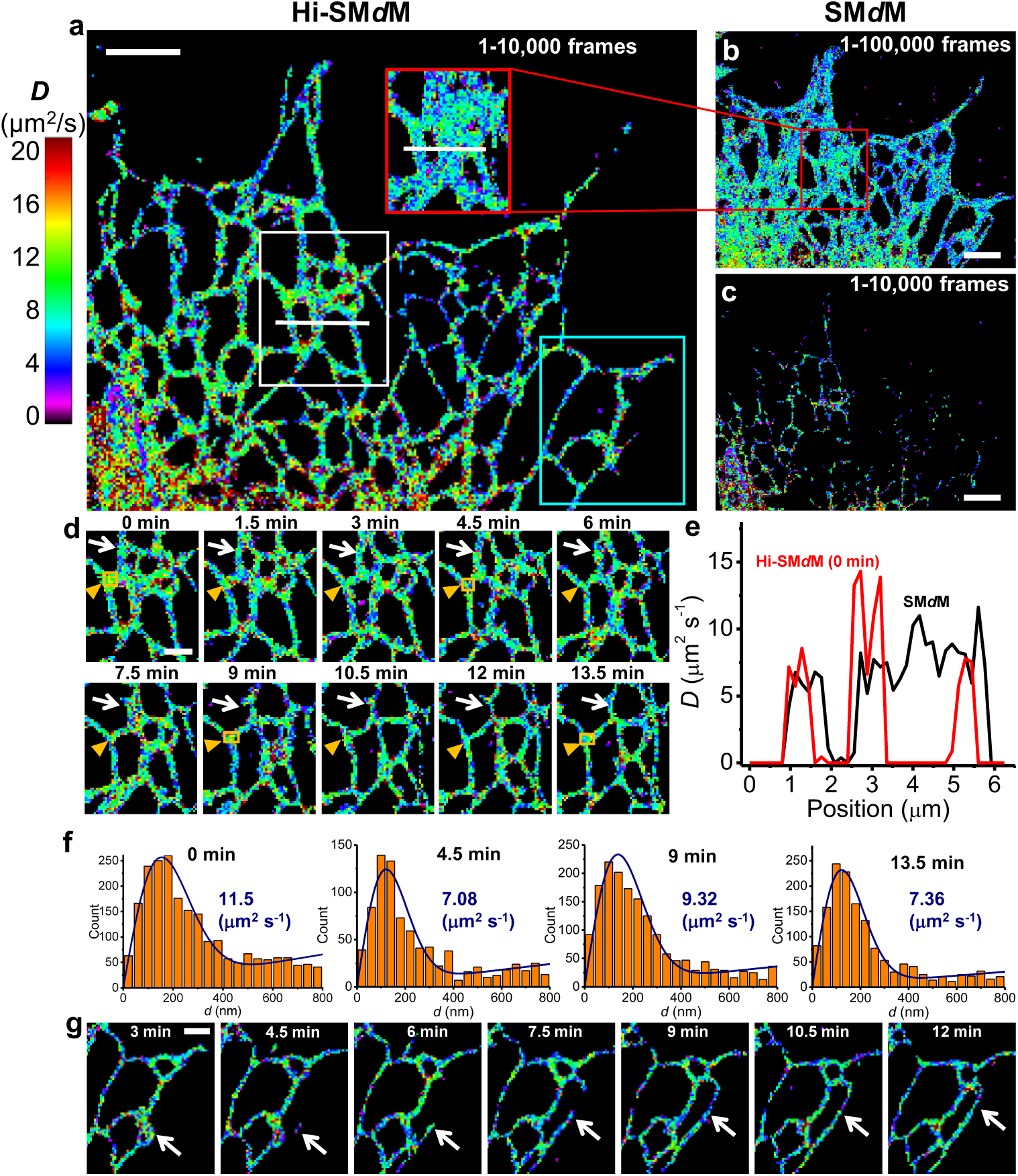
Hi-SM*d*M unveils spatiotemporal dynamics of intraorganellar diffusivity in ER. **a.** Hi-SM*d*M-reconstructed high-resolution *D* map of Dentra2 in ER lumen of a live COS-7 cell with 10,000 frames (bin size: 100×100 nm^2^). **b, c.** SM*d*M-reconstructed *D* map of Dentra2 in ER lumen of the same live COS-7 cell with 100,000 frames (b) and 10,000 frames (c). Bin size: 100×100 nm^2^. **d.** Time-resolved imaging of *D* in the boxed subregion in a (white) for every 10,000 frames at 1.5-min intervals, showing dynamic diffusivity changes in the ER lumen correlated with morphological changes in the tubule network. **e.** Variations in the *D* along the white lines in the boxed regions in a (white box) and b (red box), demonstrating that Hi-SM*d*M offers significantly higher spatial resolution compared to SM*d*M. **f.** Displacement distributions for an ER-junction area marked with orange boxes in d at different time points. Blue curves: Fits to the diffusion model, with resultant *D* values marked in each plot. **g.** Time-resolved imaging of *D* in the boxed subregion in a (cyan) at 1.5-min intervals, showing the diffusivity dynamics during the growth of new ER tubules. Scale bars: 5 μm (a, b, c); 2 μm (d, g).

Hi-SM*d*M, however, allowed continuous visualization of diffusivity changes in the ER with a moderate 1.5-minute time interval while maintaining high spatial resolution, even as the network rapidly reorganized (white arrow in Fig. 6d). This superior spatial resolution, quantified by *D* profiles extracted along the white lines in Fig. 6a, is illustrated in Fig. 6e. Diffusivity within the ER lumen fluctuated alongside tubule rearrangements (Fig. 6d, Supplementary Fig. S9 and Video 6). Quantitative analysis revealed dynamic changes in 𝐷 at the same ER tubule junction over time (orange box and arrowheads in Fig. 6d and 6f), indicating that macromolecular crowding in the ER lumen varies with morphological changes in the network. Additionally, Hi-SM*d*M captured the formation of new ER tubules, which initially exhibited slower diffusion— suggesting higher crowding—but quickly reached diffusivity levels comparable to established tubules (white arrow, Fig. 6g).

### Hi-SM*d*M unveils spatiotemporal dynamics of intraorganellar diffusivity in mitochondria

We next examined spatiotemporal dynamics of diffusion in the mitochondrial matrix. Hi-SM*d*M successfully reconstructed the intricate and clear morphology of mitochondria while minimizing motion blur by utilizing significantly fewer frames compared to previous studies^19^ (Fig. 7a and Supplementary Fig. S10). From the super-resolution *D* map, we observed nanoscale heterogeneity in diffusivity within individual mitochondria, as well as variations among mitochondria within the same cell. Notably, mitochondria near the nucleus and cell periphery exhibited higher diffusivity (Fig. 7a). Sub-micrometer patches of elevated diffusivity within a single mitochondrion likely correspond to local crista geometry. Moreover, mitochondria in specific regions, such as near the nucleus or cell periphery, may have more dynamic or open cristae in response to energy demands, resulting in higher diffusivity.

**Figure 7.**
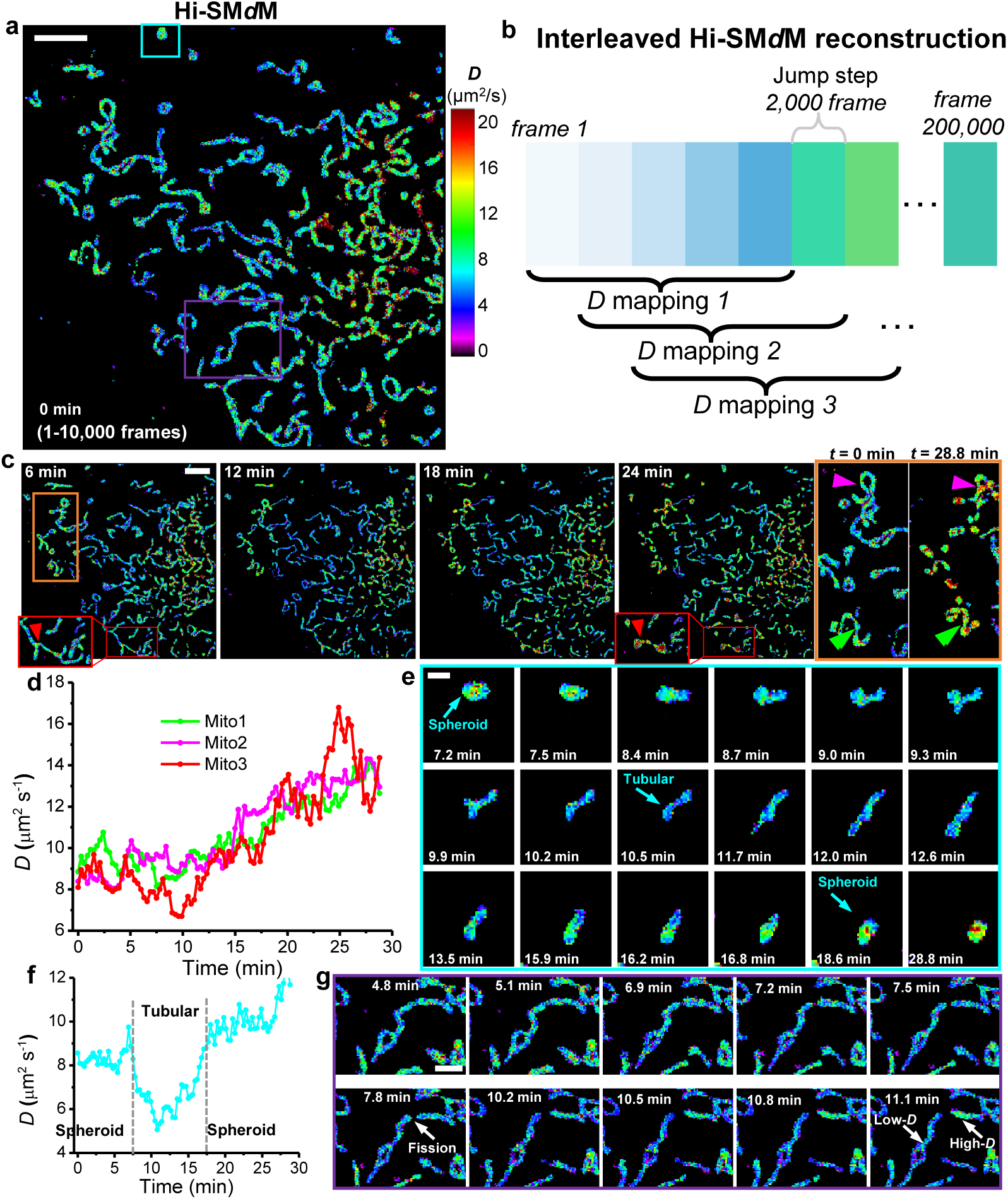
Hi-SM*d*M unveils spatiotemporal dynamics of intraorganellar diffusivity in mitochondria. **a.** Hi- SM*d*M-reconstructed high-resolution *D* map of Dentra2 in mitochondrial matrix of a live COS-7 cell with 10,000 frames (bin size: 100×100 nm^2^). **b.** Illustration of interleaved reconstruction strategies in Hi-SM*d*M by using overlapping image sets. Each color box represents 2,000-frame single-molecule image data. While conventional methods reconstruct one *D* map every 10,000 frames, Hi-SM*d*M shifts the reconstruction window by 2,000 frames to create four additional interleaved *D* maps between consecutive conventional time points. This overlapping strategy produces a series of *D* maps with a time interval of 0.3 minutes, significantly enhancing temporal resolution. **c.** Hi- SM*d*M-reconstructed full *D* maps of Dentra2 in mitochondrial matrix at 6 min, 12 min, 18 min and 24 min, revealing a progressive increase in diffusivity as mitochondria undergo gradual fragmentation (arrowheads indicated mitochondria in boxed regions). **d.** Temporal variations of *D* for the three mitochondria marked with arrowheads in c. **e.** Time-resolved imaging of *D* of the mitochondrion marked with cyan box in a, showing dynamic diffusivity changes in the matrix correlated with mitochondrial morphological changes. **f.** *D* value time traces for the same mitochondrion in e. **g.** Time-resolved imaging of *D* of the mitochondria in the boxed subregion in a (purple) at 0.3- min intervals, showing that the two mitochondria, which originated from the same mitochondrial fission, exhibit differences in diffusivity. Scale bars: 5 μm (a, c); 1 μm (e) ; 2 μm (g).

To achieve a clearer and more continuous view of dynamic changes in mitochondrial diffusivity over time, we implemented an interleaved reconstruction strategy to enhance temporal resolution. While each reconstruction still requires 10,000 frames, shifting the reconstruction window by 2,000 frames enables the incorporation of new temporal information, generating updated *D* maps as soon as data becomes available (Fig. 7b). This overlapping approach produces interleaved super-resolution *D* maps with a temporal interval of 0.3 minutes (Supplementary Video 7). Prolonged laser illumination gradually induced mitochondrial fragmentation (arrowheads in Fig. 7c). Interestingly, unlike in the cytoplasm, diffusivity within fragmented mitochondria increased (Fig. 7c, 7d and Supplementary Video 8). This effect may be attributed to the transition of mitochondria from tubular to spherical shapes and the open cristae. Mitochondria are highly dynamic organelles that undergo constant remodeling through processes like fission, fusion, and shape changes. We observed that the morphological transition between tubular and spherical mitochondria was consistently accompanied by synchronous changes in diffusivity between low and high values in real time (Fig. 7e, 7f and Supplementary Video 9). These dynamics allow mitochondria to rapidly adapt their structure to meet cellular energy demands, respond to environmental cues, and maintain mitochondrial quality. In the case of mitochondrial fission, although the overall diffusivity of the mitochondrion did not show significant changes before division, the two segments resulting from fission exhibited noticeable differences in diffusivity (arrows in Fig. 7g and Supplementary Video 10). Since diffusivity within mitochondria typically reflects the matrix structure, internal reorganization, and the presence of regulatory proteins or obstacles, these findings suggest changes in the distribution and dynamic properties of molecules. They may also indicate rapidly changing energy demands within mitochondria during or after division. Together, Hi- SM*d*M showcases exceptional temporal and spatial resolution, enabling the observation of spatiotemporal changes in diffusivity associated with highly dynamic subcellular structures and activities.

## Discussion

Mapping intracellular diffusivity is challenging, while quantifying its spatiotemporal dynamics poses an even greater difficulty. In this work, we developed a deep-learning denoising framework specifically designed for freely diffusing single-molecule images recorded under typical SM*d*M conditions with stroboscopic illumination^46^. This framework unbiasedly restores single-molecule displacements with high throughput, enabling super-resolution imaging of temporal dynamics in intracellular diffusivity. Using this approach, we uncovered rich spatiotemporal heterogeneities in molecular diffusivity within the cytoplasm and organelles of live cells.

Unlike previous deep-learning denoising methods, which relied on spatial redundancy and long-range temporal correlations to restore stationary molecules^31^, the stochastic behavior of freely diffusing single molecules disrupts these temporal correlations. To address this, we introduced Hi-SM*d*M—a denoising framework for SM*d*M that leverages noise modeling in a self-supervised context tailored for diffusing single-molecule imaging. This robust approach recovers nearly all valid molecules from noisy images while accurately preserving their diffusivity. Critically, Hi-SM*d*M addresses the inherent measurement bias of SM*d*M, where dim, fast-moving molecules with low SNR are often excluded, skewing diffusion results toward the slower-moving regime. By overcoming this bias, Hi-SM*d*M ensures high accuracy in diffusivity measurements while simultaneously achieving over 8-fold improvement in throughput. Simulations with known ground truths demonstrated Hi-SM*d*M’s ability to correctly map spatial diffusivity differences by detecting nearly all molecules and preserving their true displacements.

In live COS-7 cells, Hi-SM*d*M revealed nanoscale heterogeneities in cytoplasmic diffusivity associated with mitochondria. For the first time, it enabled visualization of dynamic structural rearrangements and mitochondrial movements by tracking temporal changes in nanoscale diffusivity. These findings surpass the capabilities of conventional techniques: FCS failed to reveal mitochondrial influences on diffusion^47^, while FRET-based biosensors could not measure crowding variations linked to mitochondria^44^. Hi- SM*d*M directly resolved the reduction in diffusion coefficients at the nanoscale and connected it to mitochondrial structures. Cytoplasmic crowding plays a crucial role in regulating protein activities and functions within cells, and molecular diffusion offers a direct means of assessing intracellular crowding. Hi-SM*d*M also demonstrated its ability to study cytoplasmic crowding under osmotic stress. By investigating live cells exposed to a 50% hypotonic environment, Hi-SM*d*M showed that macromolecular crowding decreased due to cell swelling, followed by recovery via regulatory volume increase mechanisms. With its high spatial and temporal resolution, Hi-SM*d*M is uniquely suited to uncover crowding-related processes and their functional implications across diverse cellular structures.

For the ER, Hi-SMdM provided a unique advantage. While SM*d*M could not resolve the ER’s dynamic tubule network due to motion blur from prolonged imaging, Hi-SM*d*M captured continuous diffusivity changes at 1.5-minute intervals with high spatial resolution. These observations revealed how macromolecular crowding in the ER lumen varies with morphological changes in the network, offering new insights into processes like protein synthesis, lipid metabolism, or stress responses^48^.

For the mitochondria, we implemented an interleaved reconstruction strategy to enhance temporal resolution by 5-fold and obtained a more continuous view of dynamic changes in mitochondrial diffusivity over time. We revealed the diffusivity dynamics in matrix associated with the rapid morphological changes of mitochondria and mitochondrial fission in real time. These structural transitions, particularly between tightly packed and open cristae, enable mitochondria to adapt to energy demands and stress conditions^49^. The high spatiotemporal resolution of Hi-SM*d*M offers new insights into mitochondrial energy metabolism, quality control, and the associated structural and functional dynamics.

Together, characterizing intracellular diffusivity dynamics requires live-cell imaging at both sufficient temporal and spatial resolutions to follow the fate of biological activities. Hi-SM*d*M offers a transformative approach to map diffusivity with nanoscale resolution and high accuracy, uncovering previously inaccessible heterogeneities and dynamic processes.

## Methods

### Optical setup

Single-molecule displacement/diffusivity mapping experiments, including conventional SM*d*M and Hi-SM*d*M were conducted on a Nikon Ti2-E inverted fluorescence microscope. The setup incorporated a 405 nm laser (MDL-III-405) for photoactivating mEos3.2 to its ’red’ form, a 488 nm laser (OBIS 488 LX, Coherent) for exciting the non-photoactivated ’green’ form of mEos3.2 and Dentra2, a 561 nm laser (MGL-FN-561) for exciting the photoactivated ’red’ form of mEos3.2, and a 639 nm laser (MRL-FN-639) for exciting MitoTracker. These lasers were collinearly combined and directed through a dichroic mirror (ZT405/488/561/640rpcv2) before being focused at the back focal plane of an oil-immersion objective lens (Nikon CFI Plan Apochromat λ 100×, NA 1.45). By adjusting the translation stage, the laser beams were positioned slightly below the critical angle at the sample-coverslip interface, illuminating a few micrometers into the sample with a beam diameter exceeding 50 micrometers. Fluorescence emissions excited by the 488 nm, 561 nm, and 639 nm lasers were filtered through respective emission filters (ET535/70m, ET605/70m, ET705/100m) and captured by an EMCCD camera (iXon Ultra 897, Andor). The system featured a multifunctional I/O board (PCI-7633, NI) to receive camera signals, enabling synchronization of the EMCCD exposure with stroboscopic modulation of the 561 nm laser pulses *via* an acousto-optic tunable filter (AOTF) (97-03151-01, Gooch & Housego).

### Data simulation

Single-molecule images were simulated based on two-dimensional Brownian diffusion. For the images used to train the Hi-SM*d*M network and those shown in Fig. S1, individual molecular trajectories were generated using a diffusion coefficient *D* and a simulation time step of 1 μs. Brownian motion was modeled over 1000 micro-steps, with each step representing a random displacement derived from the specified *D*, corresponding to diffusion over the duration of a laser pulse. The point spread function (PSF) for the simulations was obtained from experimental measurements of 100 nm fluorescent beads under the microscope. This PSF was applied to the simulated trajectories and projected onto a pixelated image grid, ensuring a realistic representation of single-molecule images consistent with experimental conditions.

### Training and inference

Two models were trained for noise modeling and denoising. For robust noise modeling (Fig. 2c), images acquired from various samples and fluorescence backgrounds were used. A training set of 1,000 images was employed by default, and the total loss function was optimized by combining the weighted losses from three modules: the global noise level prediction module, the local noise map prediction module, and the noise generation module (see Supplementary Note 1 for details). For the denoising network, low-SNR images (256 × 256 pixels) were used as inputs, paired with clear signal images as targets. These target images were generated under diverse simulated backgrounds to enhance generalization. Data augmentation, including random flipping and rotation, was applied to expand the dataset. Typically, 500 images were used for training the denoising network. The network optimization was guided by a second-order loss function, specifically mean squared error (MSE), defined as:

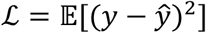

where 𝑦 represents the target signal image, and *ŷ* is the predicted denoised image. This loss function minimizes overfitting to the background, preventing excessive smoothing and ensuring the preservation of critical signal features. Both models were trained on an NVIDIA GeForce RTX 4070 GPU (12 GB), using the Adam optimizer for optimization. The learning rate was set to 0.0001, with exponential decay rates of 0.9 and 0.999 for the first and second moments, respectively. The final denoising model was achieved after 200 training epochs and demonstrated robust generalization. This strong generalization capability allowed the final trained model to be consistently applied to all experimental data throughout the study.

### Cell culture and transfection

18 mm diameter coverslips were treated in methanol solution with ultrasonic for 5min, and then rinsed with deionized water. COS-7 (Cell Resource Center, IBMS, CAMS/PUMS) cells were grown at 37 °C and 5% CO2 in Dulbecco’s modified Eagle’s medium (DMEM) (Gibco C11995500BT) supplemented with 10% fetal bovine serum (Gemini), 1× GlutaMAX Supplement, and 1× non-essential amino acids. Dendra2-ER5 was a gift from Michael Davidson (Addgene #57716). mEos3.2-C1 was a gift from Michael Davidson and Tao Xu (Addgene #54550). Mito-Dendra2 was constructed by replacing PhiYFP in the mito-PhiYFP (pPhi-Yellow-mito, Evrogen #FP607) with Dendra2 between the BamHⅠ and HindⅢ sites. The Clover-mRuby2 FRET crowding sensor was constructed in our lab and deposited on Addgene (Addgene #171058)^44^. Cells were transiently transfected with each of the above plasmids 24 hours after being plated on the precleaned glass coverslips. Transfection was performed using Lipofectamine 2000 (ThermoFisher) according to the manufacturers’ instructions. Imaging was performed 48 hours after transfection.

### Hi-SM*d*M of live cells

Hi-SM*d*M experiments in live cells were performed in DMEM (Gibco 21063029) supplemented with 25 mM HEPES at pH 7.4 (Gibco 15630106), as described previously ^45^. For a typical recorded frame size of 256×256 pixels (∼41×41 µm^2^ sample area), the EMCCD camera exposure time and dead time were 9.0 ms and 157 µs, respectively, hence camera frame time *T* = 9.16 ms, corresponding to a frame rate of 109.3 frames per second. To achieve subframe temporal resolution, paired 561 nm excitation pulses were used. One pulse was applied near the end of the odd frame, and the second pulse was positioned at the start of the even frame. The motion of single molecule was recorded over short time windows as defined by the center-to-center separation of the two pulses, Δ*t* (1 ms typical, Fig. 1 a). This paired tandem excitation scheme was repeated between 8–15 × 10^4^ times (typically), generating the final Hi- SM*d*M time-resolved image sequence (12,500 times for a single Hi-SM*d*M map of cytoplasm and 5,000 times for a single Hi-SM*d*M map of ER). For concurrent imaging of mitochondrion, MitoTracker Deep Red FM (ThermoFisher A66440) with a concentration of 30 nmol/L was added to the sample after Hi-SM*d*M imaging on stage. For the hypotonic experiment, an equal volume of water was added to the imaging medium additionally.

### Data analysis for Hi-SM*d*M

As previously described^16^, super-localization and nearest neighbor searching of molecules between tandem frames were performed after denoising, with the displacement data subsequently accumulated. The diffusion coefficients were estimated using maximum likelihood estimation (MLE) by fitting them to a probability model derived from spatially binned displacement data within a typical bin of 100×100 nm^2^ (or 160×160 nm^2^ as indicated). For the diffusivity in cytoplasm, this probability model was grounded in the principles of two-dimensional random walk, expressed as 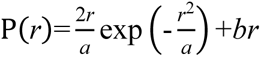. For the diffusivity in the ER lumen^19^, single-molecule displacements in the bin were next projected along the principal direction for MLE fitting to a modified one dimensional random-walk model with the probability distribution of 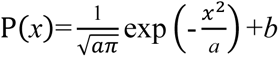. Here *a= 4DΔt*, and *b* represents a background that accounts for mismatched molecules diffusing randomly into the search radius. Performing MLE across all spatial bins yielded a spatially resolved map of local diffusion coefficients (*D*), which was subsequently utilized for color-coded rendering.

## Data availability

The data supporting this study’s findings are available from the corresponding author upon reasonable request.

## Funding

This work was supported by the National Natural Science Foundation of China (62475032, 62205048) and the Sichuan Science and Technology Program (2023NSFSC1390). J.F. acknowledges support from National Natural Science Foundation of China grant (No. 32271271) and the Sichuan Local Science and Technology Development Project guided by central government grant (No. 2024ZYD0163).

## Author Contributions

K.C. conceived the research. J.X. and Y.H. designed and conducted the experiments. Z.Z. and J.Y. built the microscope setup. All authors contributed to experimental designs, data analysis, and paper writing.

## Notes

### Competing Interest Statement

The authors have declared no competing interest.

## References

(1) Dix, J. A.; Verkman, A. Crowding effects on diffusion in solutions and cells. Annu. Rev. Biophys. 2008, 37 (1), 247–263, DOI: 10.1146/annurev.biophys.37.032807.125824

(2) Boersma, A. J.; Liu, B.; Poolman, B. A sensor for quantification of macromolecular crowding in living cells. Biophysical Journal 2015, 108 (2), 114a, DOI:10.1038/nmeth.3257

(3) Grassmann, G.; Miotto, M.; Desantis, F.; Di Rienzo, L.; Tartaglia, G. G.; Pastore, A.; Ruocco, G.; Monti, M.; Milanetti, E. Computational Approaches to Predict Protein–Protein Interactions in Crowded Cellular Environments. Chemical Reviews 2024, 124 (7), 3932–3977, DOI: 10.1021/acs.chemrev.3c00550

(4) Rivas, G.; Minton, A. P. Macromolecular crowding in vitro, in vivo, and in between. Trends in biochemical sciences 2016, 41 (11), 970–981, DOI: 10.1016/j.tibs.2016.08.013

(5) Mika, J. T.; Poolman, B. Macromolecule diffusion and confinement in prokaryotic cells. Current opinion in biotechnology 2011, 22 (1), 117–126, DOI:10.1016/j.copbio.2010.09.009

(6) Pack, C.; Saito, K.; Tamura, M.; Kinjo, M. Microenvironment and effect of energy depletion in the nucleus analyzed by mobility of multiple oligomeric EGFPs. Biophysical journal 2006, 91 (10), 3921–3936, DOI: 10.1529/biophysj.105.079467

(7) Michaluk, P.; Rusakov, D. A. Monitoring cell membrane recycling dynamics of proteins using whole-cell fluorescence recovery after photobleaching of pH-sensitive genetic tags. Nature Protocols 2022, 17 (12), 3056–3079, DOI: 10.1038/s41596-022-00732-4

(8) Lippincott-Schwartz, J.; Snapp, E. L.; Phair, R. D. The development and enhancement of FRAP as a key tool for investigating protein dynamics. Biophysical journal 2018, 115 (7), 1146–1155, DOI: 10.1016/j.bpj.2018.08.007

(9) Digman, M. A.; Gratton, E. Lessons in fluctuation correlation spectroscopy. Annual review of physical chemistry 2011, 62 (1), 645–668, DOI: 10.1146/annurev-physchem-032210-103424

(10) Krieger, J. W.; Singh, A. P.; Bag, N.; Garbe, C. S.; Saunders, T. E.; Langowski, J.; Wohland, T. Imaging fluorescence (cross-) correlation spectroscopy in live cells and organisms. Nature protocols 2015, 10 (12), 1948–1974, DOI:10.1038/nprot.2015.100

(11) Sezgin, E.; Schneider, F.; Galiani, S.; Urbanč ič, I.; Waithe, D.; Lagerholm, B. C.; Eggeling, C. Measuring nanoscale diffusion dynamics in cellular membranes with super-resolution STED–FCS. Nature protocols 2019, 14 (4), 1054–1083, DOI:10.1038/s41596-019-0127-9

(12) Manley, S.; Gillette, J. M.; Patterson, G. H.; Shroff, H.; Hess, H. F.; Betzig, E.; Lippincott-Schwartz, J. High-density mapping of single-molecule trajectories with photoactivated localization microscopy. Nature methods 2008, 5 (2), 155–157, DOI:10.1038/nmeth.1176

(13) Kusumi, A.; Tsunoyama, T. A.; Hirosawa, K. M.; Kasai, R. S.; Fujiwara, T. K. Tracking single molecules at work in living cells. Nature chemical biology 2014, 10 (7), 524–532, DOI:10.1038/nchembio.1558

(14) Cognet, L.; Leduc, C.; Lounis, B. Advances in live-cell single-particle tracking and dynamic super-resolution imaging. Current opinion in chemical biology 2014, 20, 78–85, DOI: 10.1016/j.cbpa.2014.04.015

(15) Lelek, M.; Gyparaki, M. T.; Beliu, G.; Schueder, F.; Griffié, J.; Manley, S.; Jungmann, R.; Sauer, M.; Lakadamyali, M.; Zimmer, C. Single-molecule localization microscopy. Nature reviews methods primers 2021, 1 (1), 39, DOI: 10.1038/s43586-021-00038-x

(16) Xiang, L.; Chen, K.; Yan, R.; Li, W.; Xu, K. Single-molecule displacement mapping unveils nanoscale heterogeneities in intracellular diffusivity. Nature methods 2020, 17 (5), 524–530, DOI: 10.1038/s41592-020-0793-0

(17) Xiang, L.; Chen, K.; Xu, K. Single molecules are your quanta: A bottom-up approach toward multidimensional super-resolution microscopy. ACS nano 2021, 15 (8), 12483–12496, DOI: 10.1021/acsnano.1c04708

(18) Yan, R.; Chen, K.; Xu, K. Probing nanoscale diffusional heterogeneities in cellular membranes through multidimensional single-molecule and super-resolution microscopy. Journal of the American Chemical Society 2020, 142 (44), 18866–18873, DOI:10.1021/jacs.0c08426

(19) Xiang, L.; Yan, R.; Chen, K.; Li, W.; Xu, K. Single-molecule displacement mapping unveils sign-asymmetric protein charge effects on intraorganellar diffusion. Nano letters 2023, 23 (5), 1711–1716, DOI: 10.1038/s41592-020-0793-0

(20) Choi, A. A.; Xiang, L.; Li, W.; Xu, K. Single-molecule displacement mapping indicates unhindered intracellular diffusion of small (≲ 1 kDa) solutes. Journal of the American Chemical Society 2023, 145 (15), 8510–8516, DOI: 10.1021/jacs.3c00597

(21) Li, X.; Li, Y.; Zhou, Y.; Wu, J.; Zhao, Z.; Fan, J.; Deng, F.; Wu, Z.; Xiao, G.; He, J. Real-time denoising enables high-sensitivity fluorescence time-lapse imaging beyond the shot-noise limit. Nature Biotechnology 2023, 41 (2), 282–292, DOI:10.1038/s41587-022-01450-8

(22) Speiser, A.; Müller, L.-R.; Hoess, P.; Matti, U.; Obara, C. J.; Legant, W. R.; Kreshuk, A.; Macke, J. H.; Ries, J.; Turaga, S. C. Deep learning enables fast and dense single-molecule localization with high accuracy. Nature methods 2021, 18 (9), 1082–1090, DOI: 10.1038/s41592-021-01236-x

(23) Nehme, E.; Weiss, L. E.; Michaeli, T.; Shechtman, Y. Deep-STORM: super-resolution single-molecule microscopy by deep learning. Optica 2018, 5 (4), 458–464, DOI:10.1364/OPTICA.5.000458

(24) Nehme, E.; Freedman, D.; Gordon, R.; Ferdman, B.; Weiss, L. E.; Alalouf, O.; Naor, T.; Orange, R.; Michaeli, T.; Shechtman, Y. DeepSTORM3D: dense 3D localization microscopy and PSF design by deep learning. Nature methods 2020, 17 (7), 734–740, DOI:10.1038/s41592-020-0853-5

(25) Liu, J.; Li, Y.; Chen, T.; Zhang, F.; Xu, F. Machine Learning for Single-Molecule Localization Microscopy: From Data Analysis to Quantification. Analytical Chemistry 2024, 96 (28), 11103–11114, DOI:10.1021/acs.analchem.3c05857

(26) Weigert, M.; Schmidt, U.; Boothe, T.; Müller, A.; Dibrov, A.; Jain, A.; Wilhelm, B.; Schmidt, D.; Broaddus, C.; Culley, S. Content-aware image restoration: pushing the limits of fluorescence microscopy. Nature methods 2018, 15 (12), 1090–1097, DOI:10.1038/s41592-018-0216-7

(27) Chaudhary, S.; Moon, S.; Lu, H. Fast, efficient, and accurate neuro-imaging denoising via supervised deep learning. Nature communications 2022, 13 (1), 5165, DOI:10.1038/s41467-022-32886-w

(28) Li, X.; Zhang, G.; Wu, J.; Zhang, Y.; Zhao, Z.; Lin, X.; Qiao, H.; Xie, H.; Wang, H.; Fang, L. Reinforcing neuron extraction and spike inference in calcium imaging using deep self-supervised denoising. Nature methods 2021, 18 (11), 1395–1400, DOI:10.1038/s41592-021-01225-0

(29) Lequyer, J.; Philip, R.; Sharma, A.; Hsu, W.-H.; Pelletier, L. A fast blind zero-shot denoiser. Nature Machine Intelligence 2022, 4 (11), 953–963, DOI:10.1038/s42256-022-00547-8

(30) Zhang, G.; Li, X.; Zhang, Y.; Han, X.; Li, X.; Yu, J.; Liu, B.; Wu, J.; Yu, L.; Dai, Q. Bio-friendly long-term subcellular dynamic recording by self-supervised image enhancement microscopy. Nature Methods 2023, 20 (12), 1957–1970, DOI:10.1038/s41592-023-02058-9

(31) Li, X.; Hu, X.; Chen, X.; Fan, J.; Zhao, Z.; Wu, J.; Wang, H.; Dai, Q. Spatial redundancy transformer for self-supervised fluorescence image denoising. Nature Computational Science 2023, 3 (12), 1067–1080, DOI 10.1038/s43588-023-00568-2

(32) Abdelhamed, A.; Brubaker, M. A.; Brown, M. S. Noise flow: Noise modeling with conditional normalizing flows. In Proceedings of the IEEE/CVF International Conference on Computer Vision, 2019; pp 3165–3173, DOI:10.1109/ICCV.2019.00326

(33) Brooks, T.; Mildenhall, B.; Xue, T.; Chen, J.; Sharlet, D.; Barron, J. T. Unprocessing images for learned raw denoising. In Proceedings of the IEEE/CVF conference on computer vision and pattern recognition, 2019; pp 11036–11045, DOI: 10.1109/CVPR.2019.01129

(34) Guo, L.; Huang, S.; Liu, H.; Wen, B. Towards robust image denoising via flow-based joint image and noise model. IEEE Transactions on Circuits and Systems for Video Technology 2023, DOI: 10.1109/TCSVT.2023.3345667

(35) Guo, S.; Yan, Z.; Zhang, K.; Zuo, W.; Zhang, L. Toward convolutional blind denoising of real photographs. In Proceedings of the IEEE/CVF conference on computer vision and pattern recognition, 2019; pp 1712–1722, DOI: 10.1109/CVPR.2019.00181

(36) Jang, G.; Lee, W.; Son, S.; Lee, K. M. C2n: Practical generative noise modeling for real-world denoising. In Proceedings of the IEEE/CVF International Conference on Computer Vision, 2021; pp 2350–2359, DOI: 10.1109/ICCV48922.2021.00235

(37) Fu, Z.; Guo, L.; Wen, B. sRGB Real Noise Synthesizing with Neighboring Correlation-Aware Noise Model. In Proceedings of the IEEE/CVF Conference on Computer Vision and Pattern Recognition, 2023; pp 1683–1691, DOI: 10.1109/CVPR52729.2023.00168

(38) Nam, S.; Hwang, Y.; Matsushita, Y.; Kim, S. J. A holistic approach to cross-channel image noise modeling and its application to image denoising. In Proceedings of the IEEE conference on computer vision and pattern recognition, 2016; pp 1683–1691, DOI: 10.1109/CVPR.2016.186

(39) Ronneberger, O.; Fischer, P.; Brox, T. U-net: Convolutional networks for biomedical image segmentation. In Medical image computing and computer-assisted intervention–MICCAI 2015: 18th international conference, Munich, Germany, October 5-9, 2015, proceedings, part III 18, 2015; Springer: pp 234-241, DOI:10.1007/978-3-319-24574-4_28

(40) Zhang, M.; Chang, H.; Zhang, Y.; Yu, J.; Wu, L.; Ji, W.; Chen, J.; Liu, B.; Lu, J.; Liu, Y. Rational design of true monomeric and bright photoactivatable fluorescent proteins. Nature methods 2012, 9 (7), 727–729, DOI:10.1038/nmeth.2021

(41) Kuimova, M. K.; Botchway, S. W.; Parker, A. W.; Balaz, M.; Collins, H. A.; Anderson, H. L.; Suhling, K.; Ogilby, P. R. Imaging intracellular viscosity of a single cell during photoinduced cell death. Nature chemistry 2009, 1 (1), 69–73, DOI: 10.1038/nchem.120

(42) Śmigiel, W. M.; Mantovanelli, L.; Linnik, D. S.; Punter, M.; Silberberg, J.; Xiang, L.; Xu, K.; Poolman, B. Protein diffusion in Escherichia coli cytoplasm scales with the mass of the complexes and is location dependent. Science Advances 2022, 8 (31), eabo5387, DOI: 10.1126/sciadv.abo5387

(43) Guo, M.; Pegoraro, A. F.; Mao, A.; Zhou, E. H.; Arany, P. R.; Han, Y.; Burnette, D. T.; Jensen, M. H.; Kasza, K. E.; Moore, J. R. Cell volume change through water efflux impacts cell stiffness and stem cell fate. Proceedings of the National Academy of Sciences 2017, 114 (41), E8618–E8627, DOI: 10.1073/pnas.1705179114

(44) Chen, K.; Yan, R.; Xiang, L.; Xu, K. Excitation spectral microscopy for highly multiplexed fluorescence imaging and quantitative biosensing. Light: Science & Applications 2021, 10 (1), 97, DOI: 10.1038/s41377-021-00536-3

(45) Yan, J.; He, Y.; Chen, K. Excitation Spectral Microscopy for Highly Multiplexed Intracellular Imaging via Quasi-Blind Unmixing. ACS Photonics 2024, 11 (8), 3454–3466, DOI:10.1021/acsphotonics.4c01037.

(46) Elf, J.; Li, G.-W.; Xie, X. S. Probing transcription factor dynamics at the single-molecule level in a living cell. Science 2007, 316 (5828), 1191–1194, DOI: 10.1126/science.1141967

(47) Baum, M.; Erdel, F.; Wachsmuth, M.; Rippe, K. Retrieving the intracellular topology from multi-scale protein mobility mapping in living cells. Nature communications 2014, 5 (1), 4494, DOI:10.1038/ncomms5494

(48) Monterroso, B.; Margolin, W.; Boersma, A. J.; Rivas, G.; Poolman, B.; Zorrilla, S. Macromolecular crowding, phase separation, and homeostasis in the orchestration of bacterial cellular functions. Chemical reviews 2024, 124 (4), 1899–1949,DOI: 10.1021/acs.chemrev.3c00622

(49) Ježek, P.; Jabů rek, M.; Holendov á, B.; Engstov á, H.; Dlaskov á, A. Mitochondrial cristae morphology reflecting metabolism, superoxide formation, redox homeostasis, and pathology. Antioxidants & Redox Signaling 2023, 39 (10), 635–683, DOI: 10.1089/ars.2022.0173

